# Hippocampal fear engrams modulate ethanol-induced maladaptive contextual generalization in mice

**DOI:** 10.1101/2022.01.24.477538

**Authors:** C. Cincotta, E. Ruesch, R. Senne, S. Ramirez

**Affiliations:** Department of Psychological and Brain Sciences, Boston University, Boston MA, 02215

**Keywords:** addiction, alcohol, withdrawal, fear conditioning, extinction, engram, hippocampus, optogenetics

## Abstract

The compounding symptomatology of comorbid alcohol use disorder (AUD) and post-traumatic stress disorder (PTSD) gives rise to an interaction of maladaptive neurobiological processes, the etiology of which remains elusive. Here, we devised an optogenetic strategy aimed at rescuing maladaptive responses to fearful stimuli in male c57BL/6 mice following chronic ethanol administration and forced abstinence. In the first experiment, we confirmed that fear acquisition and maladaptive contextual generalization was potentiated in ethanol-exposed mice during fear conditioning and exposure to a novel environment, respectively. In the second experiment, using an activity-dependent tet-tag system, we labeled and artificially inhibited the neural ensemble selectively activated by contextual fear conditioning in the dorsal hippocampus to attenuate behavioral dysfunctions resulting from ethanol exposure. We found that acute optogenetic inhibition during exposure to a novel environment suppressed maladaptive generalization in ethanol-exposed mice. These results provide further evidence for a crucial link between ethanol exposure and impaired fear memory processing by providing cellular and behavioral insights into the neural circuitry underlying AUD and PTSD comorbidity.

## MAIN TEXT

The comorbidity of alcohol use disorder (AUD) and post-traumatic stress disorder (PTSD) poses a unique challenge to society. Epidemiological evidence of co-occurring PTSD and AUD spans decades and continents, revealing the vast global impact of these disorders with one another (Helzer et al., 1987; Bremner et al., 1996; Mills et al., 2006; Kachadourian & Petrakis, 2016; Smith & Cottler, 2018). Dual-diagnosed individuals with PTSD and AUD experience increased negative affective mood and behaviors, as each disorder acts as a risk factor for the other (Carlson & Weiner, 2021). Markedly, research has shown that treatment options for individuals with a dual-diagnosis are less effective and have poorer outcomes compared to treating individuals diagnosed with either PTSD or AUD alone (McCauley et al., 2012). Despite the significant societal impact, the neurobiological understanding of AUD/PTSD comorbidity is not well understood, and there remains a need to investigate the neural substrates of this dual-diagnosis.

AUD and PTSD converge on similar behavioral phenomena, including but not limited to maladaptive stress responses. Recent work has sought to investigate hypotheses that these disorders converge not only on behavior, but also neural substrates and pathways (Enman et al., 2014, Gilpin & Weiner, 2017). Alcohol use disorder is a potent risk factor thought to precipitate a medley of cellular, behavioral, and cognitive pathologies that converge on memory impairments associated with hippocampal dysfunction (Goode & Maren 2019). Additionally, the role of the hippocampus in contextual fear memory acquisition and generalization has been well documented (Hainmueller & Bartos, 2020). Researchers have found that fear memories can generalize immediately following memory consolidation (Asok et al., 2019; Yu et al., 2021). Notably, contextual discrimination and generalization are impaired in both AUD and PTSD (Scarlata et al., 2019; Gilpin & Weiner, 2017).

The ability to disambiguate similar events is an integral process supporting episodic memory. Termed pattern separation, this theoretical construct posits the creation of orthogonal representations which transform similar inputs into dissimilar outputs, preventing catastrophic interference between new and old memories (Yassa & Stark., 2012). The hippocampus is a causal node for the encoding, storage, and retrieval of episodic memories (Hainmueller & Bartos, 2020). However, empirical evidence supporting the hippocampus’s role in pattern separation was largely unfounded until recently (McHugh et al., 2007; Nakazawa et al., 2004). Previous research has suggested that this cognitive process may be largely predicated on the dentate gyrus (DG) subregion of the hippocampus. (Kesner, 2018; Treves & Rolls, 1994). Experimental evidence has shown ablating the DG is sufficient to disrupt a rodent’s ability to discriminate between two spatial locations (Gilbert et al., 2001). Subtle environmental changes result in widely divergent neural representations via the remapping of place cell firing rates in the DG (Leutgeb et al., 2007). Although these findings have helped to identify the neural mechanisms underlying pattern separation, prior methodological constraints did not permit the perturbation of neuronal activity in the DG at a population level.

Promisingly, recent methodological advances provide the opportunity for further experimentation focused on uncovering how mechanisms contributing to pattern separation are preserved or modified at higher levels of neurobiological organization. The combination of optogenetic and activity-dependent strategies has recently permitted causally probing the circuits and systems involved in learning and memory (Deisseroth, 2014). In recent years, one of the primary focuses of learning and memory research has centered on engrams – the specific ensembles of cells active during memory formation. These collections of cells are the putative physical correlates of memory in the brain (Josselyn & Tonegawa, 2020). Engrams are known to undergo plasticity, and there has been substantial evidence that optogenetic activation of these ensembles is sufficient to facilitate retrieval and elicit behavioral responses (Chen et al., 2019; Liu et al., 2012; Ramirez et al., 2015). Moreover, previous work from our lab has demonstrated that stimulating putative hippocampal engrams is sufficient to induce acute and enduring changes in cellular activity and behavior (Ramirez et al. 2015, Chen et al. 2019). Recently, our lab’s investigation of maladaptive behavioral responses resulting from comorbid AUD and PTSD showed that chronic optogenetic activation of engrams in the dorsal dentate gyrus (dDG) of the hippocampus was sufficient to reduce fear responses in ethanol-exposed mice (Cincotta et al. 2021). Therefore, we asked if artificial suppression of a hippocampal contextual-fear engram could mitigate ethanol-induced maladaptive generalization to better understand how this mechanism is impaired in animals displaying addiction-related behaviors.

We first utilized a behavioral strategy to evaluate maladaptive contextual generalization in a mouse model of chronic ethanol exposure and forced abstinence. All subjects were treated in accordance with protocol 201800579 approved by the Institutional Animal Care and Use Committee at Boston University. In the first experiment, thirty-two adult male mice were placed on a diet containing doxycycline (40 mg/kg; Dox). Briefly, mice were injected with 250 nL of a virus cocktail containing pAAV9-cFos-tTA and pAAV9-TRE-eYFP into the dDG at a rate of 100 nL/min (Figure 1a, see also Chen et al., 2019; Ramirez et., 2015). Following recovery, mice received 5 consecutive days of either ethanol (EtOH) or saline (Sal) treatment via intraperitoneal injection (30% ethanol (vol/vol) in saline (0.9%) at a dose of 2.0 g/kg), followed by a 48 hours forced abstinence period (Cincotta et al., 2021; Pina & Cunningham, 2017; Quinones-Laracuente et al., 2015), which coincided with the removal of Dox from their diet to allow tagging during fear conditioning (Figure 1b).

**Figure 1.**
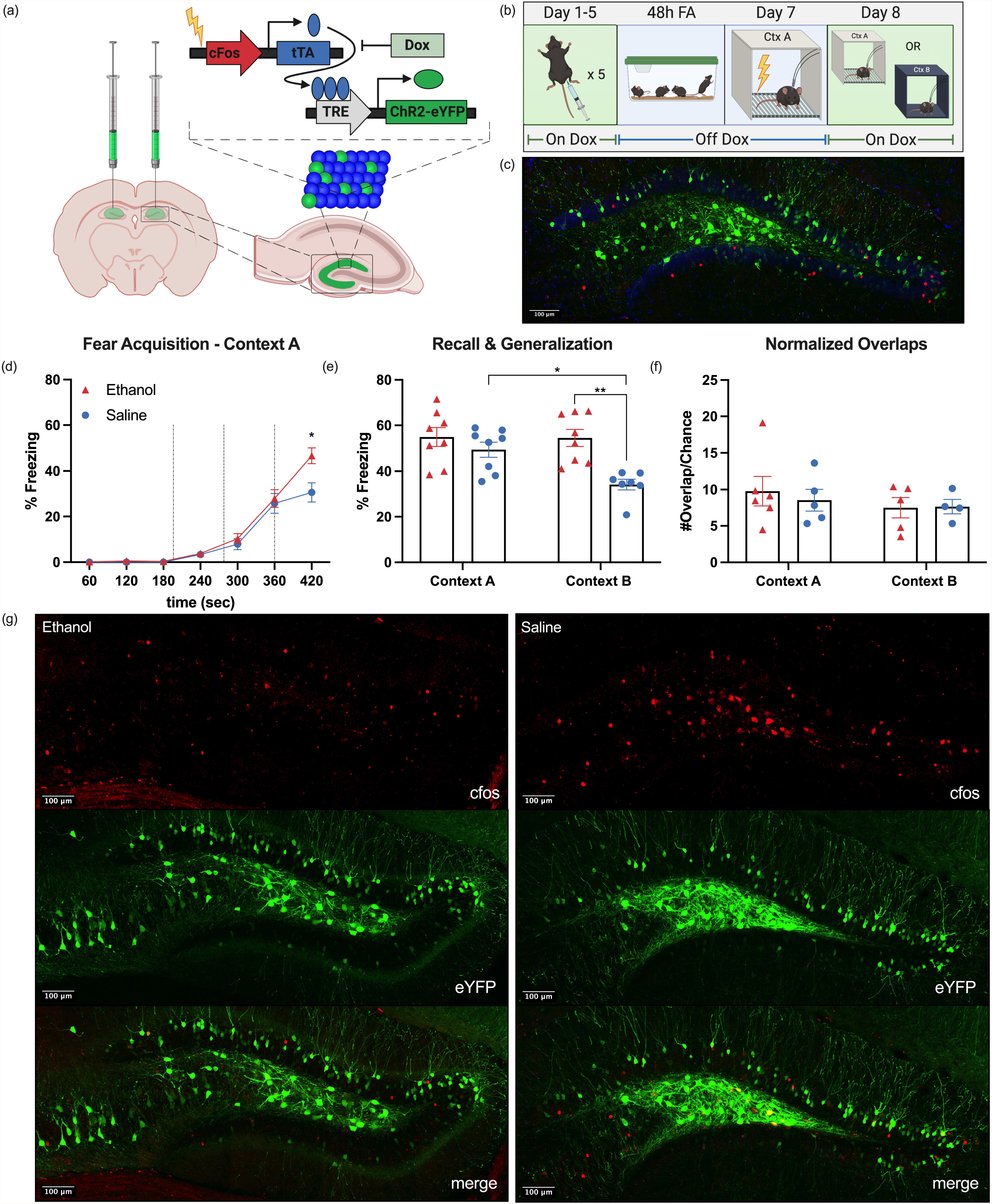
Ethanol-induced maladaptive contextual generalization. (a) Schematic of viral strategy. A viral cocktail of AAV9-c-Fos-tTA and AAV9-TRE-eYFP was infused into the dorsal dentate gyrus (dDG) for activity-dependent transcription of eYFP. Schematic created with BioRender. (b) Behavioral design for ethanol (EtOH) or saline exposure, forced abstinence, fear conditioning, and recall or generalization tests. Schematic created with BioRender. (c) Representative histology for activity-dependent tagging of contextual fear engrams. Scale bar = 100 μm. (d) Fear acquisition in Context A. No group differences in freezing during fear acquisition were observed (two-way RM ANOVA, treatment F_(1,174)_= 3.648, P=0.0661). Main effect of time (two-way RM ANOVA, time F_(6,174)_=99.15, P<0.001). Time x Treatment interaction (two-way RM ANOVA, time x treatment F_(6,174)_=3.440, P=0.0031). (e) Recall or generalization tests in neutral Context B. Main effects and interaction of context and treatment (two-way ANOVA, treatment F_(1,27)_= 13.51, P=0.0010, context F_(1,27)_= 4.907, P=0.0354, context x treatment F_(1,27)_= 4.416, P=0.0451). EtOH-exposed mice froze significantly higher than the saline control mice in novel Context B (Tukey’s HSD p=0.0023). Saline-control mice froze significantly less in Context B relative to Context A (Tukey’s HSD p=0.0276), while the EtOH-exposed mice showed no difference across contexts (Tukey’s HSD p=0.9998). (f) Quantified normalized overlaps. See main text for normalization process. No significant differences across treatment groups or contexts in terms of percent of reactivated cells from the tagged putative engram (two-way ANOVA, context F_(1,16)_=0.8946, p=0.3585, treatment F_(1,16)_=0.1045, p=0.7507). (g) Representative histology for dual-labelled engram cells reactivated during recall or generalization. Scale bar = 100 μm.

All mice were fear conditioned in Context A (Ctx A); Room A, Coulbourne mouse chambers (insert dimensions), grid flooring, sanitized with 70% EtOH, dimmed white lighting, and handled by experimenter 1. Following a 198s baseline period, mice received three mild shocks, 2 second each at 0.5 mA, with an ITI of 78s and 60s following the last shock. No group differences in freezing during fear acquisition were observed (Figure 1d) (two-way RM ANOVA, treatment F_(1,174)_= 3.648, P=0.0661), however we observed a main effect of time (two-way RM ANOVA, time F_(6,174)_=99.15, P<0.001) and a time x treatment interaction (two-way RM ANOVA, time x treatment F_(6,174)_=3.440, P=0.0031), demonstrating learning and acquisition of fear. At the last time point, Sidak’s post-hoc comparison revealed EtOH-exposed mice were freezing at a significantly higher level than saline control mice, indicating that EtOH exposure may result in heightened fear responses during acquisition as the quantity of shocks increased.

The following day, mice were either placed back into Ctx A for a 5-minute recall test, or into a novel Context B (Ctx B) for a 5-minute generalization test (Ctx B; Room B, red lighting, Coulbourne mouse chambers, vertical black and white stripes on front and back chamber panels, solid black covering on side panels, grid flooring covered with semi-transparent plastic sheets, scented with almond-extract, sanitized with Rescue Disinfectant, and handled by experimenter 2).

Across testing, we found significant main effects of both context and treatment, as well as a treatment x context interaction (Figure 1e) (two-way ANOVA, treatment F_(1,27)_= 13.51, P=0.0010, context F_(1,27)_= 4.907, P=0.0354, context x treatment F_(1,27)_= 4.416, P=0.0451). As expected, Tukey’s HSD revealed no group differences between EtOH-exposed and saline control mice in Ctx A (p=0.6729). Consistent with previous studies, EtOH-exposed mice froze significantly higher than the saline control mice in novel Context B (Tukey’s HSD p=0.0023) (Sarlata et al., 2019). Additionally, the saline control mice froze significantly less in Context B relative to Context A (Tukey’s HSD p=0.0276), while the EtOH-exposed mice showed no difference across contexts (Tukey’s HSD p=0.9998).

All mice were perfused 90-minutes following either recall or generalization tests to capture peak cFos expression. Tissue samples were treated and stained as in Chen et al. 2019. eYFP/ArchT+ cells were quantified using Ilastik, a machine-learning based bioimage analysis pipeline, using the pre-defined pixel classification and object segmentation workflows (Berg et al. 2019). Randomly chosen representative histology was input into Ilastik as training data for the respective algorithms. cFos+ cells and dual labeled “overlap” cells were identified and counted by hand using ImageJ’s CellCounter, with the experimenter blind to treatment and context groups. Due to visual differences in expression of eYFP and ArchT, it was not possible to blind the experimenter to this variable. Once the total number of cFos, eYFP/ArchT, and “overlap” cells were determined, the relative amount of each cell type was normalized to the total number of DAPI cells present in each given slice of the dDG (cFos/DAPI, eYFP or ArchT/DAPI, and overlap/DAPI). Total number of DAPI cells were estimated by first quantifying a sample of representative slices via Ilastik processing, and then determining the average DAPI per area in order to apply this average ratio to the area of all slices quantified for histology. Chance of overlap was defined as (cFos/DAPI) * (eYFP or ArchT/DAPI). Overlap/Chance determined the normalized overlap per slice, which was averaged among all slices per animal to calculate the normalized percentage of overlap between cFos positive cells and the tagged engram. Despite behavioral differences in freezing across contexts, we found no significant differences across treatment groups or contexts in terms of the percent of cells reactivated from the tagged putative engram (Figure 1f) (two-way ANOVA, context F_(1,16)_=0.8946, p=0.3585, treatment F_(1,16)_=0.1045, p=0.7507). See representative histology (Figure 1g).

Previous work from our lab showed the effectiveness of using optogenetic activation to modulate addiction-related states in EtOH-exposed mice (Cincotta et al., 2021). Therefore, we asked if optogenetic perturbation of the putative hippocampal engram could similarly mediate the EtOH-induced contextual overgeneralization observed in the first experiment. To do so, 72 male mice were surgerized as previously described with a virus cocktail containing pAAV9-cFos-tTA and pAAV9-TRE-ArchT-eYFP (i.e. an inhibitory opsin) or with pAAV9-TRE-eYFP as a control (Figure 2a). All mice were treated with either ethanol or saline as described in experiment one. Following fear conditioning in Context A, mice underwent either a 5-minute recall or generalization test in Context A or Context B, respectively (Figure 2b). Coinciding with the testing period, mice receive 5 minutes of optogenetic inhibition of the dDG cells tagged previously in Context A (15 ms pulse width, 20 Hz).

**Figure 2.**
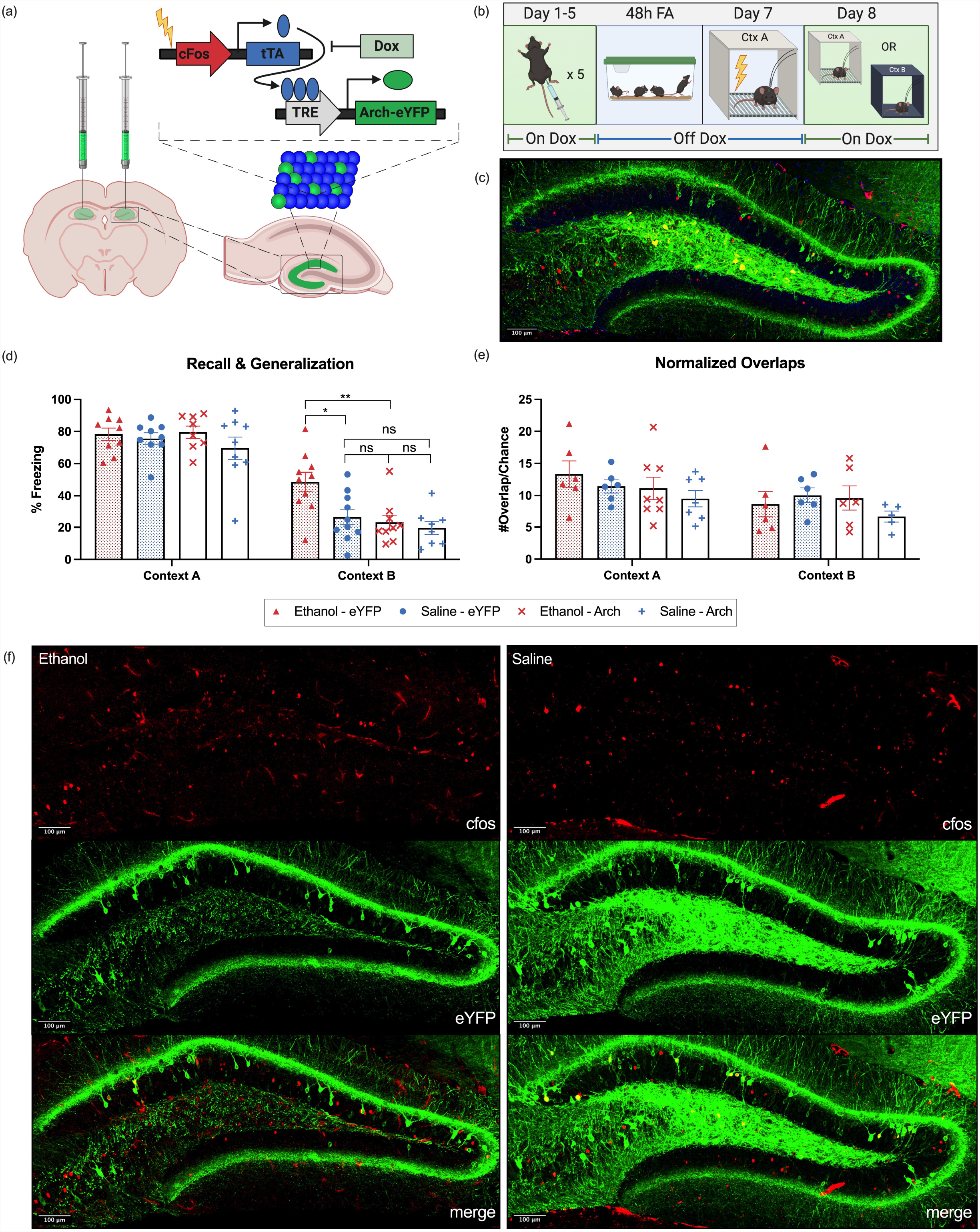
ArchT inhibition of a contextual fear memory prevents ethanol-induced maladaptive generalization. (a) Schematic of viral strategy. A viral cocktail of AAV9-c-Fos-tTA and AAV9-TRE-ArchT or AAV9-TRE-eYFP was infused into the dorsal dentate gyrus (dDG) for activity-dependent transcription of ArchT or eYFP. Schematic created with BioRender. (b) Behavioral design for ethanol (EtOH) or saline exposure, forced abstinence, fear conditioning, and recall or generalization tests. Schematic created with BioRender. (c) Representative histology for activity- dependent tagging of contextual fear engrams. Scale bar = 100 μm. (d) Optogenetic inhibition during recall or generalization tests. Main effects of context, virus, and treatment (three-way ANOVA, context F_(1,64)_=174.4, p<0.0001; virus F_(1, 64)_=6.963, p=0.0104; treatment F_(1,64)_=7.3, p=0.0085). Strong trending interactions of context x virus (F_(1,64)_=3.860, p=0.0538), and context x virus x treatment (F_(1,64)_=3.428, p=0.0687). No significant difference in freezing between ArchT EtOH-exposed mice and ArchT saline-control mice in Context B (Tukey’s post-hoc p=0.9997), and ArchT EtOH-exposed mice froze at a significantly lower level than the eYFP EtOH-exposed mice in Context B (Tukey’s post-hoc, p=0.0089), indicating optogenetic inhibition was able to significantly decrease freezing in EtOH-exposed mice, preventing mice from overgeneralizing to the neutral context. (e) Quantified normalized overlaps. See main text for normalization process. Analysis revealed a main effect of context (three-way ANOVA, Context F_(1, 42)_=5.180, p=0.0280). No significant interactions or individual group differences. (f) Representative histology for dual-labelled engram cells reactivated during recall or generalization. Scale bar = 100 μm.

Analysis of freezing behavior of all mice during the optogenetic inhibition revealed a main effect of context, virus, and treatment (three-way ANOVA, context F_(1,64)_=174.4, p<0.0001; virus F_(1, 64)_=6.963, p=0.0104; treatment F_(1,64)_=7.3, p=0.0085). We observed strong, albeit not significant, interactions of context x virus (F_(1,64)_=3.860, p=0.0538), and context x virus x treatment (F_(1,64)_=3.428, p=0.0687) (Figure 2d). Post-hoc analysis of individual group differences confirmed that there was no significant difference across all groups in Context A - all mice, regardless of treatment or virus group, expressed high levels of freezing during the recall session, with no effect of optogenetic inhibition on freezing. Excitingly, however, in Context B, we observed that optogenetic inhibition of tagged dDG cells was able to significantly decrease freezing in EtOH-exposed mice, preventing mice from overgeneralizing to the neutral context. As in experiment one, the eYFP EtOH-exposed mice froze significantly higher than the eYFP saline-control mice in context B (Tukey’s post-hoc p=0.0292). However, there was no significant difference in freezing between ArchT EtOH-exposed mice and ArchT saline-control mice in Context B (Tukey’s post-hoc p=0.9997). The ArchT EtOH-exposed mice froze at a significantly lower level than the eYFP EtOH-exposed mice in Context B (Tukey’s post-hoc, p=0.0089), indicating that optogenetic inhibition was successful in reducing aberrant fear responses in a neutral context.

As in the first experiment, mice were perfused 90-minutes following behavior and tissue samples were treated for histology accordingly. Our analysis revealed only a main effect of context (three-way ANOVA, Context F_(1, 42)_=5.180, p=0.0280) with no significant interactions or individual group differences. Altogether, these data suggest that despite similar levels of cFos activation and putative engram reactivation across groups, perturbation of the dDG is sufficient to mitigate addiction-related states and prevent mice from expressing EtOH-induced overgeneralization in a neutral context.

In light of research examining how rodents withdrawn from EtOH exhibit impairments in fear memory extinction and maladaptive generalization (Cincotta et al., 2021; Scarlata et al., 2019) here we sought to investigate how EtOH-exposure and withdrawal would affect rodents during a contextual fear conditioning. We first predicted that cessation of EtOH administration would cause overgeneralization of fear to the neutral environment, and that these deficits could be rescued by optogenetic inhibition of the contextual-fear engram stored in the dDG. As expected, mice that had been previously treated with alcohol exhibited a tendency to freeze at higher levels than saline controls during fear acquisition. This result dovetails with previous work showing how EtOH withdrawal is sufficient to potentiate conditioned fear learning (Bertotto et al., 2006). Interestingly, we also found that EtOH exposure and forced abstinence facilitated the generalization of fear between the conditioned context and a novel neutral context, demonstrating that EtOH withdrawal can induce maladaptive generalization of traumatic experiences. In our paradigm, we believe that forced abstinence of ethanol precipitated acute withdrawal-induced changes to the stress and memory systems in our mice, which led to the maladaptive overgeneralization seen in a neutral context. Taken together, these results provide further evidence that EtOH dependence and subsequent withdrawal causes dysfunction to the neural circuitry underlying emotional stress that results in potentiated behavioral responses to stressors.

Our previous work demonstrated that chronic optogenetic stimulation of tagged dDG cells is sufficient to ameliorate EtOH-induced extinction recall impairments (Cincotta et al., 2021) and that acute optogenetic activation of a fear memory can transiently restore freezing (Ramirez et al., 2013). Accordingly, we posited that acute optogenetic inhibition of the dDG-mediated contextual fear engram would sufficiently attenuate EtOH-induced deficits in contextual generalization.

Initially, we found mice that underwent ethanol exposure, forced abstinence, and contextual fear conditioning (in Ctx A), exhibited heightened levels of freezing in a neutral environment (Ctx B) relative to saline-control animals indicating EtOH-induced maladaptive generalization (Figure 1). However, when cells active during the fear conditioning session were tagged with an inhibitory opsin, acute optogenetic inhibition of the putative engram was sufficient to eliminate disparities in neutral context generalization between treatment groups (Figure 2). Interestingly, optogenetic inhibition of the tagged cells during a recall session in the fear conditioning context elicited no changes in freezing relative to eYFP-control groups, suggesting the effect seen in the neutral context was not the result of an overall reduction in fear.

We speculate that inhibiting the hippocampal node responsible for encoding the contextual elements of a fear memory engram was sufficient to bypass the dysregulation of numerous brain-wide systems implicated in mediating addiction-related behaviors, such as the insular cortex, which regulates interoceptive feelings of drug craving and withdrawal (Naqvi and Bechara., 2009). At the molecular level, EtOH administration has also been shown to aberrantly affect hippocampal neurogenesis (Golub et al., 2015) and neuroinflammation (Tajuddin et al., 2014). The dDG has been proposed to be responsible for decreasing memory interference by promoting pattern separation, i.e. the process by which overlapping hippocampal inputs are orthogonalized into more dissimilar outputs, thereby decreasing interference between similar memories (Favila et al., 2016). Consequently, our optogenetic perturbation may be artificially facilitating more precise hippocampal indexing by silencing the competing fear memory trace to effectively prevent destructive interference and transiently pausing the pro-inflammatory and aberrant neurogenesis resulting from EtOH exposure, which future work may address directly.

Additionally, we found no difference in cFos+ cell quantifications between EtOH-exposed and Sal-control groups under both normative and optogenetic conditions. Although this finding may be constrained by the size of our treatment groups, it is more likely that an upstream brain region is mediating the behavioral impairment observed in normative EtOH-exposed mice. For instance, previous studies examining how EtOH alters fear-memory retrieval found increased cFos+ expression in the prelimbic cortex, paraventricular thalamus, and medial central amygdala, and have found these regions function as a circuit during fear retrieval (Quinones-Laracuente et al., 2015; Do-Monte et al., 2015). Future work may quantify overlaps in these regions and test if they are more predictive of behavioral discrimination in comparison to the dDG. Indeed, the emergence of novel strategies capable of mitigating fear responses holds promising value in better understanding the underlying mechanisms of learning and memory alongside psychiatric disorders including PTSD. The tagging and artificial perturbation of engrams enables us to catalogue the neurobiological changes resulting from addiction-related pathologies (Whitaker and Hope., 2018). By further characterizing the cellular, circuit-level, and systems-wide modifications that arise during states of drug- or alcohol-dependence, we hope to facilitate the development of novel therapeutic interventions aimed at artificially and enduringly restoring healthy neuronal function.

## Acknowledgements

This work was supported by a Ludwig Family Foundation grant, an NIH Early Independence Award (DP5 OD023106-01), an NIH Transformative R01 Award, a Young Investigator Grant from the Brain and Behavior Research Foundation, the McKnight Foundation Memory and Cognitive Disorders award, the Center for Systems Neuroscience and Neurophotonics Center at Boston University. The authors would also like to thank Cornwall’s in Kenmore Square, for providing a supportive environment and space in which this manuscript was written.

## Declaration of Interests

The authors declare no competing interests.

## Notes

### Competing Interest Statement

The authors have declared no competing interest.

